# Rapid response of nitrogen cycling gene transcription to labile carbon amendments in a soil microbial community

**DOI:** 10.1101/2021.03.26.437204

**Authors:** Peter F. Chuckran, Viacheslav Fofanov, Bruce A. Hungate, Ember M Morrissey, Egbert Schwartz, Jeth Walkup, Paul Dijkstra

## Abstract

Episodic inputs of labile carbon (C) to soil can rapidly stimulate nitrogen (N) immobilization by soil microorganisms. However, the transcriptional patterns that underlie this process remain unclear. In order to better understand the regulation of N cycling in soil microbial communities, we conducted a 48 h laboratory incubation with an agricultural soil where we stimulated the uptake of inorganic N by amending the soil with glucose. We analyzed the metagenome and metatranscriptome of the microbial communities at four timepoints that corresponded with changes in N availability. The relative abundances of genes remained largely unchanged throughout the incubation. In contrast, glucose addition rapidly increased transcription of genes encoding for ammonium and nitrate transporters, enzymes responsible for N assimilation into biomass, and genes associated with the N regulatory network. This upregulation coincided with an increase in transcripts associated with glucose breakdown and oxoglutarate production, demonstrating a connection between C and N metabolism. When concentrations of ammonium were low, we observed a transient upregulation of genes associated with the nitrogen fixing enzyme nitrogenase. Transcripts for nitrification and denitrification were downregulated throughout the incubation, suggesting that dissimilatory transformations of N may be suppressed in response to labile C inputs in these soils. These results demonstrate that soil microbial communities can respond rapidly to changes in C availability by drastically altering the transcription of N cycling genes.

**IMPORTANCE:** A large portion of activity in soil microbial communities occurs in short time frames in response to an increase in C availability, affecting the biogeochemical cycling of nitrogen. These changes are of particular importance as nitrogen represents both a limiting nutrient for terrestrial plants as well as a potential pollutant. However, we lack a full understanding of the short-term effects of labile carbon inputs on the metabolism of microbes living in soil. Here, we found that soil microbial communities responded to labile carbon addition by rapidly transcribing genes encoding proteins and enzymes responsible for inorganic nitrogen acquisition, including nitrogen fixation. This work demonstrates that soil microbial communities respond within hours to carbon inputs through altered gene expression. These insights are essential for improved understanding of the microbial processes governing soil organic matter production, decomposition, and nutrient cycling in natural and agricultural ecosystems.

## INTRODUCTION

Inorganic nitrogen (N) availability in soil dictates several ecosystem-level processes such as plant growth (1), greenhouse gas emissions in the form of nitrous oxide (2), and eutrophication from runoff (3). The transformation of N by soil microbial communities is directly tied to the pool of bioavailable N in soils (4, 5). Thus, understanding the controls of N metabolism in soil microbes is key to determining, and potentially managing (6), the cycling of N in soils. Although genes and regulatory mechanisms for microbial N cycling processes have long-been identified in laboratory studies (7–9), the short-term dynamics and controls of N cycling in complex soil communities remain poorly understood. The availability of shotgun sequencing technologies to analyze microbial functioning in soil communities provides an opportunity to enhance our understanding of microbially mediated soil N cycling.

Measuring short-term responses of soil microbial populations to changes in the environment is crucial in understanding the role of microbes in biogeochemical cycling. Most biogeochemical transformations occur during short periods of intense microbial activity, when the active fraction of microbes may be up to 20 times higher than in bulk soil (10). This stimulation is often the result of a localized increase in nutrient concentrations, such as in the rhizosphere or an area of fresh organic matter decomposition. Despite the importance of these “hot moments”, only a few studies (e.g. 11, 12) have tracked changes in N-cycling gene transcription in soils.

Notably, the short-term (hours to days) transcriptional response of N-cycling genes in response to labile C inputs has yet to be determined. Microbial communities experience sudden changes in C and N availability associated with plant root exudation (13), trophic interactions (14, 15), and litter leachate (16). Since soil microbes are typically limited by labile C and energy (17–19), the addition of a C-rich substrate is expected to stimulate growth and activity (20), increasing the demand for N (21). Whether N is derived from the uptake of organic N present in the substrate or mineral N available in the soil depends largely on the C:N of the substrate (22). For example, in Yang et al. 2016 (23) soil microbial communities assimilated organic N during the mineralization of added glycine, but in the presence of glucose the mineralization of glycine was initially suppressed and ammonium served as the main source of N. Simple sugars such as glucose have accordingly been shown to influence protease activity (24). The metabolic pathways for N immobilization have been well characterized *in vitro* (25). A majority of N assimilation into biomass occurs through the conversion of NH_4_^+^ into the amino acids glutamine and glutamate, which are used as sources of N for all other amino acids. Under low-to-moderate intracellular concentrations of NH_4_^+^, the enzymes glutamine synthetase (GS; encoded by *glnA*) and glutamate synthase (GOGAT; *gltS*) convert NH_4_^+^ to glutamate in a two-step reaction referred to as the GS-GOGAT pathway (26). Under high concentrations of NH_4_^+^, the enzyme glutamate dehydrogenase (GDH; *gudB, gdhA*) converts NH_4_^+^ directly to glutamate in a one-step reversible reaction (27).

Since both the GS-GOGAT pathway and GDH require N as NH_4_^+^, other forms of inorganic N must be converted to ammonium before conversion into biomass. In the case of nitrate and nitrite, the reduction to ammonium occurs through either assimilatory nitrate reduction or, under anoxic conditions, dissimilatory nitrate reduction to ammonium (DNRA; Table S1) (28). The conversion of atmospheric N_2_ to ammonium by diazotrophs is catalyzed by the enzyme nitrogenase (*nifDHK*) (29).

The mechanisms regulating N uptake in response to C have been extensively studied *in vitro* (8, 25). The complex regulatory network includes a specialized sigma factor (σ^54^; *rpoN*), three transcriptional regulators, and a phosphorylation cascade comprised of post-modification enzymes, PII proteins, and a two-component regulator (30). The activity of many of the enzymes and proteins in the phosphorylation cascade is tightly controlled by cellular concentrations of glutamine and oxoglutarate (31). Since the concentration of oxoglutarate is impacted by the activity of the TCA cycle, the regulation of N cycling is directly tied to C metabolism (32).

Carbon substrate addition is also thought to influence dissimilatory N cycling processes such as nitrification and denitrification. In nitrification, ammonia is oxidized to nitrite and then nitrate. Often the steps of this process occur in different organisms (33), however complete ammonia oxidizers have also been described (34, 35).”. In denitrification, nitrate is reduced to nitrite, nitric oxide, and then nitrous oxide and N_2_. Nitrification and denitrification, beyond their ability to draw from the pools of ammonium and nitrate, also represent important avenues of inorganic N loss from soils via nitrate leaching and the release of N_2_ and nitrous oxide, a potent greenhouse gas (36). The addition of glucose is expected to have both positive and negative effects on nitrification. Rates of autotrophic nitrification tend to decrease as heterotrophs outcompete autotrophic nitrifiers for ammonium (37), but rates of heterotrophic nitrification may increase after labile C inputs (38). Denitrification is more directly influenced by C availability and quality (39), and the abundance of mRNA transcripts associated with denitrification was stimulated with the addition of glucose in anoxic soil microcosms (40).

Despite our knowledge of the mechanisms and controls of N cycling and N metabolism, we do not yet fully understand how these genes are regulated within complex soil microbial communities. Metatranscriptomics allows us to capture the transcriptional profile of a microbial community, providing insight into the potential activity of a community at a given moment in time (41–43). Many studies utilizing this technique have focused on the influence of ecosystem level characteristics/properties on transcription, such as land-use, above ground cover, seasonality, and climate (e.g. 38–43). Although these studies contribute greatly to our understanding of community gene transcription, there is additional need to study the dynamic short-term responses of microbial communities to changes in C and N availability (50).

In order to fill this knowledge gap, we conducted a soil incubation study where we induced rapid immobilization of inorganic N by adding glucose. We selected glucose as it is a form of labile C commonly found in soils (51), and has been widely used to alleviate C limitation in soil microbial communities as a means to study growth (52, 53) and metabolic activity (50). We analyzed metagenomes and metatranscriptomes of the soil microbial community using high throughput shotgun sequencing to identify the response of N cycling genes over a 48-hour period. We hypothesized that the abundance of N-cycling genes in the metagenomes would not significantly change throughout the course of the 48-hour incubation, but that changes in activity would be immediately detected in the metatranscriptomes. We further hypothesized that there would be an upregulation of genes associated with inorganic N transport, N assimilation into biomass, and N metabolism regulation in response to labile C inputs, and that the abundance of these transcripts would track the concentrations of inorganic N. This work provides an in-depth look at the short-term transcriptional response of soil microbial communities during a central biogeochemical process in soils.

## METHODS

### Soil Sampling and Site Description

Soils were collected in the fall of 2017 from a long-term crop rotation experiment at the West Virginia University Certified Organic Farm near Morgantown, West Virginia, USA (39.647502° N, 79.93691° W; 243.8 – 475.2 m a.s.l.) (54, 55). Samples were taken from plots subject to a four-year conventionally tilled crop cycle consisting of corn, soybean, wheat and a mix of kale and cowpea. Manure was added every two years (during corn and wheat planting), and rye-vetch was added as a winter cover crop before replanting corn in the spring. From each plot, 10 cores 0-10 cm depth were collected and pooled.

### Laboratory Incubation

Soil samples were shipped on ice to Northern Arizona University in Flagstaff, Arizona, USA. Soils from 3 plots were pooled, cleaned of roots and large debris, passed through a 2 mm sieve, and distributed between 64 glass Mason jars (500 mL), generating microcosms containing 30 g of soil each. The soil was preincubated at lab temperature (~ 23 °C) for 2 weeks prior to the glucose addition.

The microcosms received 1.6 mL of 0.13 M glucose solution, which added 0.7 mg of glucose C g^−1^ dry soil and raised the moisture content to 60% water holding capacity. Concentrations of glucose in this range have been demonstrated to stimulate soil microbial communities without creating a detrimental increase in osmotic pressure (52). Moreover, a brief trial incubation was conducted to ensure that this concentration of glucose would stimulate CO_2_ production. Soils were incubated at lab temperature (~ 23 °C) under ambient lighting, but never direct sunlight. Every 4 hours, over a 48 h period, 5 jars were randomly selected and destructively sampled. From each jar, we measured headspace CO_2_ concentration, concentrations of NO_3_^−^ and NH_4_^+^, and microbial biomass. A portion of each sample was immediately frozen using liquid N_2_ and stored at −80°C for DNA and RNA extraction.

Since the addition of water may stimulate community activity and respiration, especially when starting with very dry soil (56, 57), we measured respiration in a parallel incubation wherein the same volume of water was added without glucose. Headspace CO_2_ from these jars was measured and compared against the glucose additions in order to determine the overall effect of glucose and water on microbial respiration.

### Biogeochemical Measurements and Analysis

To measure soil NO_3_^−^ and NH_4_^+^ concentration 8 g of soil from each destructively sampled jar were added to 40 ml of 1 M KCl solution, shaken for 1 hour, and filtered through Whatman no. 1 filter paper. Extracts were analyzed on a SmartChem 200 Discrete Analyzer (Westco Scientific Instruments, Brookfield, Connecticut, USA). Microbial biomass was measured using an extraction-fumigation-extraction technique (58), consisting of a 0.5 M K_2_SO_4_ extraction followed by a subsequent K_2_SO_4_ extraction with the addition of chloroform. The first extraction provided an estimate of the K_2_SO_4_ extractable organic C and N from each sample, while the second extraction provided an estimate of microbial biomass C (MBC) and N (MBN). Concentrations of extractable organic C and N were measured on a TOC-L (Shimadzu Corp, Kyoto, Japan). The concentration of CO_2_ from the headspace of each microcosm was measured using a LI-6262 CO_2_/H_2_O Analyzer (Licor Industries, Omaha, Nebraska, USA) as described in Dijkstra et al. (2011) (59).

### DNA and RNA Extraction and Sequencing

We extracted DNA and RNA just before (t_0_) and 8 (t_8_), 24 (t_24_), and 48 (t_48_) h after glucose addition (n=4). DNA and RNA were extracted using the RNeasy Powersoil Total RNA Kit (Qiagen) according to manufacturer instructions. DNA was separated from RNA using the RNeasy PowerSoil DNA Elution Kit (Qiagen). RNA samples were treated with RNase-Free DNase Set (Qiagen) to remove any DNA. Nucleic acid concentrations were determined with a Qubit fluorometer (Invitrogen, Carlsbad, California, USA), and purity was assessed with a NanoDrop ND-1000 spectrophotometer (Nanodrop Technologies, Wilmington, Delaware, USA). High-quality samples were sent to the Joint Genome Institute (JGI) for sequencing (60). Paired-end, 2 × 151 bp, libraries were prepared using the Illumina NovaSeq platform (Illumina Inc., San Diego, California, USA). Raw sequence reads were uploaded to the JGI genome portal (https://genome.jgi.doe.gov/portal/) under GOLD project ID Gs0135756. A more detailed description of the sequencing can be found in the data release (61).

### Metagenome and Metatranscriptomic Analysis

Metatransciptomes were assembled by JGI using MEGAHIT v1.1.2 (62) (parameters “megahit ––k–list 23,43,63,83,103,123 ––continue –o out.megahit”) and metagenomes were assembled using SPAdes version 3.13.0 (63). Assembled metatranscriptomes and metagenomes were uploaded to the Integrated Microbial Genomes and Microbiomes (IMG/M) (64) pipeline for annotation. Full details of the bioinformatics pipeline, as well as SRA reference numbers can be found in the data release (61). From IMG/M we retrieved the number of reads for all genes attributed to functional orthologs in the Kyoto Encyclopedia of Genes and Genomes (KEGG) Orthology database (65), as well as taxonomic annotations against the IMG database. Contigs are available through the JGI genome portal, and taxonomic and functional annotations of these contigs are available on the IMG/M database (http://img.jgi.doe.gov), under GOLD project ID Gs0135756. JGI Genome ID’s for each sample, as well as sample metadata, can be found in Chuckran et al (2020; 61).

Normalization of KEGG functional annotations was preformed using the Bioconductor (66) program DESeq2 (67) in R. DESeq2 uses a negative binomial distribution to normalize read counts and estimates average log_2_ fold change (LFC) between harvests. Significant LFCs for each KEGG functional gene and transcript were determined through both a likelihood ratio test (for overall significance) and Wald test (for specific contrasts between timepoints) provided in DESeq2. Significance for both tests were assumed as a false discover rate (FDR) < 0.01. Prior to analysis, genes with less than 60 reads summed over all samples were discarded in an effort to reduce the FDR correction and improve detection of significant LFCs (68).

To assess differences in genes and transcripts composition over time, we performed permutational multivariate analysis of variance (PERMANOVA) on our metagenomes and metatranscriptomes. PERMANOVAs were conducted using Bray-Curtis dissimilarity matrices of the square root transformed normalized read counts with 999 permutations. A SIMPER analysis was used to determine genes which most strongly influenced differences between harvests. PERMANOVAs and SIMPER analyses were conducted using the vegan package (69) for R.

To assess the response of N metabolism to the addition of glucose, KEGG Orthology identifiers (K numbers) were grouped according to KEGG pathways and modules associated with N cycling (70), and K numbers representing regulatory genes controlling N metabolism were identified (8, 25) (Table S1). The response of C metabolism was determined by grouping K numbers by KEGG modules associated with glucose uptake, specifically the Entner-Doudoroff pathway (KEGG module M0008), TCA cycle (M00009), pentose phosphate pathway (M00004), gluconeogenesis (M00003), and Glycolysis (M00001). From the TCA cycle, we also determined the response of isocitrate dehydrogenase (*icd*), which produces oxoglutarate, an important metabolite linking C and N metabolism (32). Counts and LFCs for K numbers were then averaged for each module to assess the overall response for each process. Results were visualized using the ggplot2 package (71) in R v 3.6.1 (72).

## RESULTS

### Biogeochemical Measurements

The concentration of NO_3_^−^ decreased in the 24 hours after glucose addition and remained low for the remainder of the incubation (Fig. 1A). The concentration of NH_4_^+^ also decreased during the first 24 hours of the incubation and increased thereafter (Fig. 1B). Rates of CO_2_ production increased from 4-16 hours and then decreased from 28-48 hours in response to glucose (Fig. 1C). We found that the addition of water only slightly influenced CO_2_ production (Fig. S1), indicating that the majority of the stimulation was due to the addition of labile C. K_2_SO_4_ extractable organic carbon decreased for the first 20 hours and plateaued thereafter (Fig. 1D). Based on these biogeochemical measurements, we selected 4 timepoints (t_0_, t_8_, t_24_, and t_48_) from which we extracted DNA and RNA. These timepoints captured distinct phases of C and N availability that enabled us to test our hypotheses.

**Figure 1.**
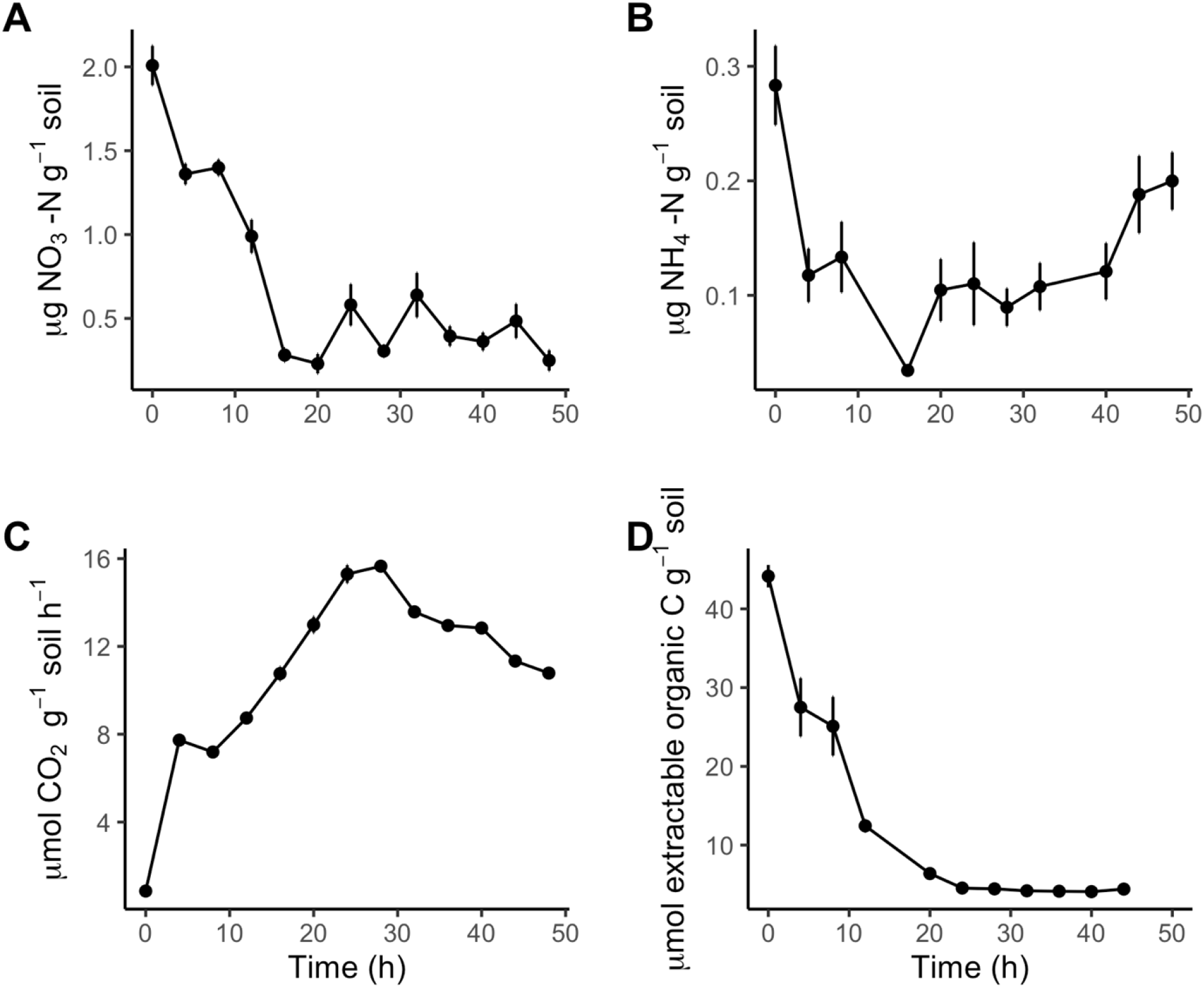
Mean concentration (± SE) of nitrate (**A),.** ammonium **(B)**, rate of carbon dioxide production **(C)**, and K_2_SO_4_-extractable C **(D)** as a function of time after glucose amendments.

Microbial biomass C (MBC) moderately decreased throughout the incubation (Fig. S2A) and microbial biomass N (MBN) remained constant (Fig. S2B). Bacteria may exhibit some stoichiometric plasticity in response to nutrient inputs (73), however a decrease in biomass C:N in response to C inputs is counter-intuitive. Since the method of microbial biomass extraction used involves two extractions on the same sample (one before and after fumigation), incomplete extraction of the added glucose in the first extraction could result in an artificially high estimate of biomass C. We believe that it is far more likely that microbial biomass and stoichiometry did not change, and that changes in estimated MBC are likely the result of unextracted glucose remaining from the initial K_2_SO_4_ extraction.

### Metagenomic and Metatranscriptomic Assembly and Annotation

Out of 16 soil samples from which DNA and RNA were extracted, 12 were successfully sequenced and assembled for metagenomic analysis and all 16 for metatranscriptomic analysis. For the metagenomes, the proportion of genes successfully annotated against the KEGG database varied from 23.4% to 25.6% of all genes per sample. Of the 6,876 functional KEGG orthologs identified in the metagenome analysis, 671 genes were in higher abundance while 332 were present in lower abundance (FDR < 0.01) after the addition of glucose. Glucose caused a shift in the relative abundance of functional genes (PERMANOVA, F_3.11_ = 3.24, P < 0.01; Fig. 2A). The genes that were most different in gene abundance relative to t_0_ varied for each timepoint (SIMPER analysis; Table S3A), and not one of these genes was directly related to N uptake. Among these were the subunits of RNA polymerase *rpoB* and *rpoC*, which were in slightly lower abundance at t_8_ (LFC −0.1, FDR > 0.1), and the regulatory gene for the lac operon, *lacI*, which was in a greater abundance at t_24_ and t_48_ (LFC 0.7, FDR < 0.01). The largest changes were found at t_24_ for low-abundant spore gemination proteins (Table S3B), specifically *gerKC* (KO6297) and *yfkQ* (K06307) which were 8.8 and 7.4 LFC more abundant than at t_0_.

**Figure 2.**
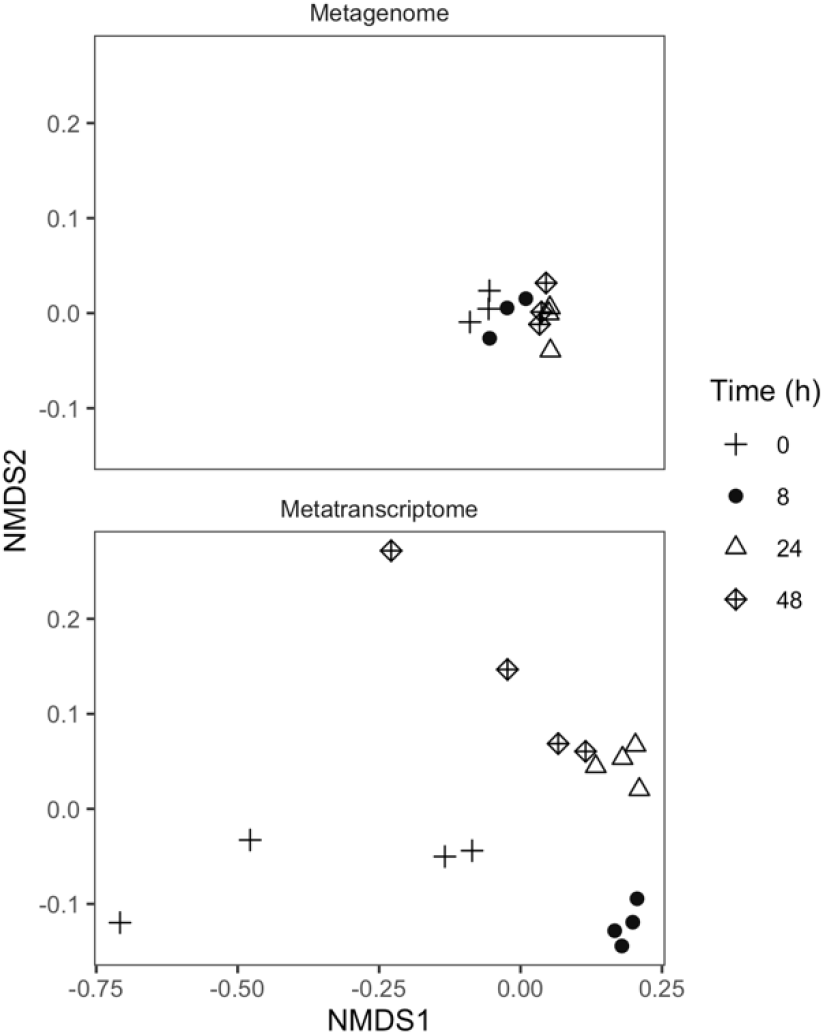
NMDS using Bray-Curtis distance of normalized KEGG annotation abundance for metagenomes (**A**) and metatranscriptomes (**B**) at 0, 8, 24, and 48 hours after the addition of glucose.

The proportion of transcripts successfully annotated against the KEGG database varied between 12.6% and 32% of all transcripts in a metatranscriptome. Transcripts for 5,448 functional genes were identified, of which 1,141 increased and 855 decreased in response to glucose. A PERMANOVA indicated significant shifts in the abundance of transcripts between timepoints (F_3, 15_ = 8.07, P < 0.01; Fig. 2B). Transcripts encoding for *amt* and *glnA* contributed the most to dissimilarity with t_0_ (SIMPER analysis), combined they explained 1% of the differences at t_8_, 1% of differences at t_24_, and 0.9% of differences at t_48_.

### Gene and Transcript Abundance of Nitrogen Cycling Processes

The abundance of N cycling genes was generally stable over time (Fig. 3A), with changes in gene abundance often being several orders of magnitude smaller than changes in transcript abundances. For metatranscriptomes, many genes associated with N uptake were highly upregulated in response to glucose (Fig. 3). Expression of genes encoding the GS-GOGAT pathway (GS - *glnA;* GOGAT *-gltS, gltD, gltB*) was consistently upregulated after glucose addition (FDR < 0.01), peaking at 8 h (Fig. 3B, Table S2). We did not find a similar trend for transcripts associated with glutamate dehydrogenase (GDH: *gudB, gdhA*). Instead we found variable increases and decreases in the expression for these genes which corresponded with different classes of GDH enzymes (Fig. 3B Table S2). In prokaryotes, GDH often uses NADH (EC 1.4.1.2), NADPH (EC 1.4.1.4) as cofactors, while GDH in eukaryotes can use both (NAD(P)H; EC 1.4.1.3) (74). Transcription of genes for EC 1.4.1.4 significantly increased early (t_8_, LFC 1.542 ± 0.312; FDR < 0.01), and transcription for EC 1.4.1.2 trended higher later (t_48_, LFC 2.229 ± 0.884; FDR < 0.1). The eukaryotic EC 1.4.1.2 gene GDH2 (K15371) was upregulated at t_24_ (LFC 1.350 ± 0.434; Table S2; FDR < 0.01) and EC 1.4.1.3 was slightly downregulated throughout (significantly at t_8_, FDR < 0.01).

**Figure 3.**
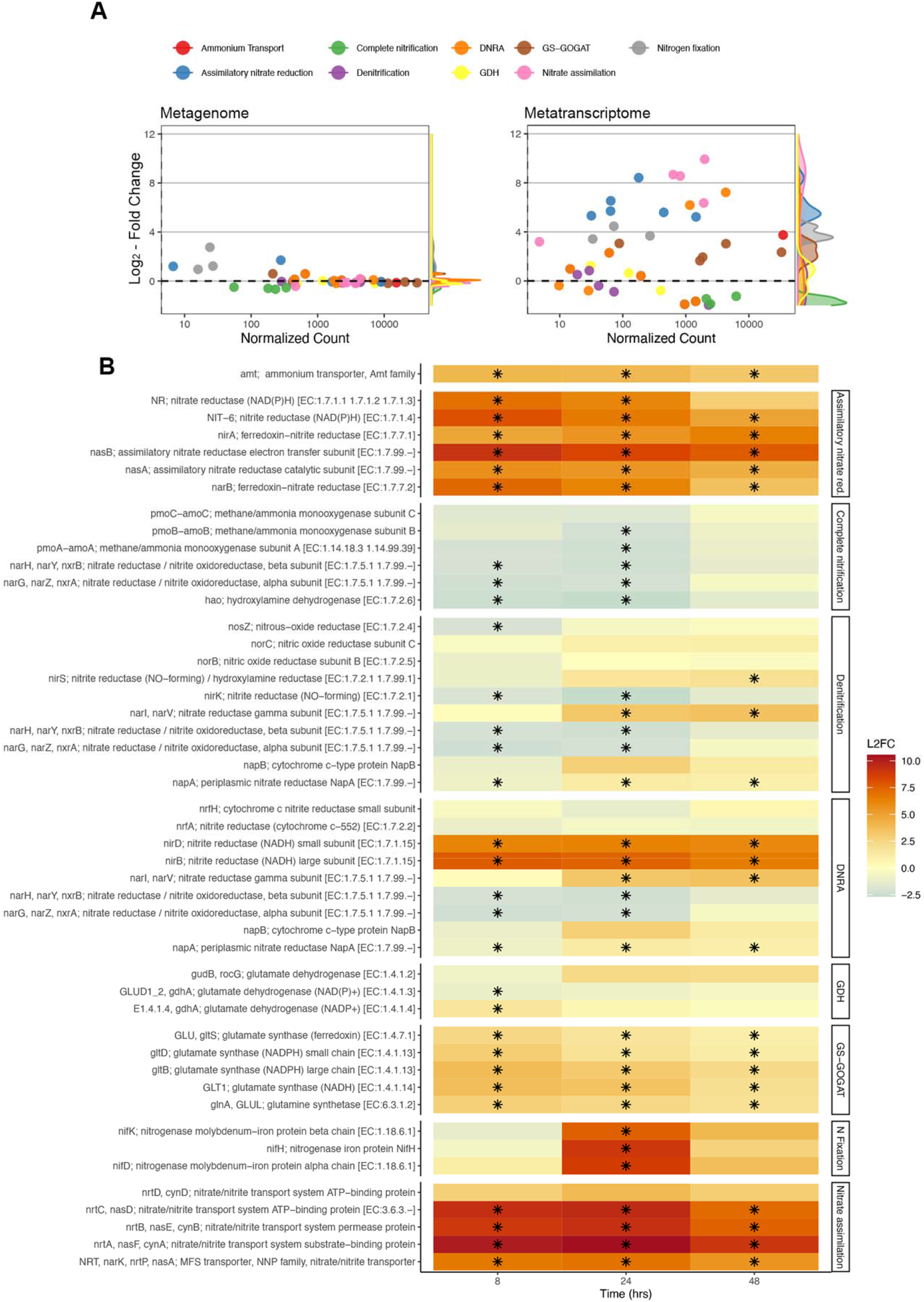
**(A)** Log_2_-fold changes (mean LFC ± SE) relative to t_0_ of normalized gene (left) and transcript (right) abundances versus normalized counts for N cycling genes from glucose-amended soils. LFC and normalized counts represent the average between t_8_, t_24_, and t_48_ for each gene. **(B)** Log_2_-fold changes in transcript abundances for genes grouped by biologically relevant reactions and pathways. A black asterisk indicates a significant change relative to t_0_.

The abundance of transcripts encoding the ammonium transporter AmtB (*amt*) was significantly (FDR < 0.01) higher after glucose addition throughout the 48-h incubation (Fig. 3B, Table S2), peaking at t_8_, where it was 16-fold higher than at t_0_ (41,366 transcripts at t_8_ vs 2,539 at t_0_). A similar upregulation was found for genes associated with nitrate and nitrite transport across the membrane – 1500-fold increases compared to t_0_ (from 2.6 to almost 2800 transcripts per sample at t_24_; Fig. 3B).

Genes associated with assimilatory nitrate reduction (Fig. 3; Table S2) were strongly upregulated at t_8_ and remained upregulated over the 48 h incubation period. In contrast, we found variable responses of genes associated with DNRA. Most genes associated with the dissimilatory reduction of nitrate to nitrite were downregulated or not significantly affected, with a few exceptions. Nitrate reductase gamma subunits (*narI/narV*) were upregulated at t_24_ and t_48_, and the genes *nirB* and *nirD*, which encode the small and large subunit of the cytosolic enzyme nitrite reductase, were significantly (FDR < 0.01) upregulated throughout the incubation (LFC 6.18 to 7.70; Fig. 3B). In contrast to these enzymes, abundance of transcripts that encode a periplasmic cytochrome c nitrite reductase (*nrfA* and *nrfH*) did not significantly change in response to C amendment.

Expression of all genes involved with nitrification were downregulated in response to glucose, and a majority of those genes (5 of 6) were significantly (FDR < 0.01) downregulated at some point during the incubation (Fig. 3B). Similarly, expression for most denitrification genes were downregulated throughout the incubation, with the exception of *narI* and *narV*, which encode for gamma subunits of nitrate reductase.

Transcripts for three genes that encode subunits of nitrogenase (*nifK, nifD,* and *nifH*) were detected, all of which were at very low abundance at t_0_, t_8_, and t_48_. Only at t_24_ did we observe a strong significant (FDR < 0.01) upregulation for all 3 genes, up to 410-fold higher than t_0_ for *nifH* (798 transcripts at t_24_ vs 1 at t_0_; Fig. 3B).

We found that the vast majority of N cycling gene transcription could be attributed to bacteria and archaea (Fig. 4). Dissimilatory processes were largely from *Thaumarchaeota* and *Nitrospirae*, while assimilatory processes tended to be represented by *Proteobacteria, Actinobateria, and Acidobacteria*. Nitrogen fixation was heavily dominated by *Proteobacteria* (Fig. 4).

**Figure 4.**
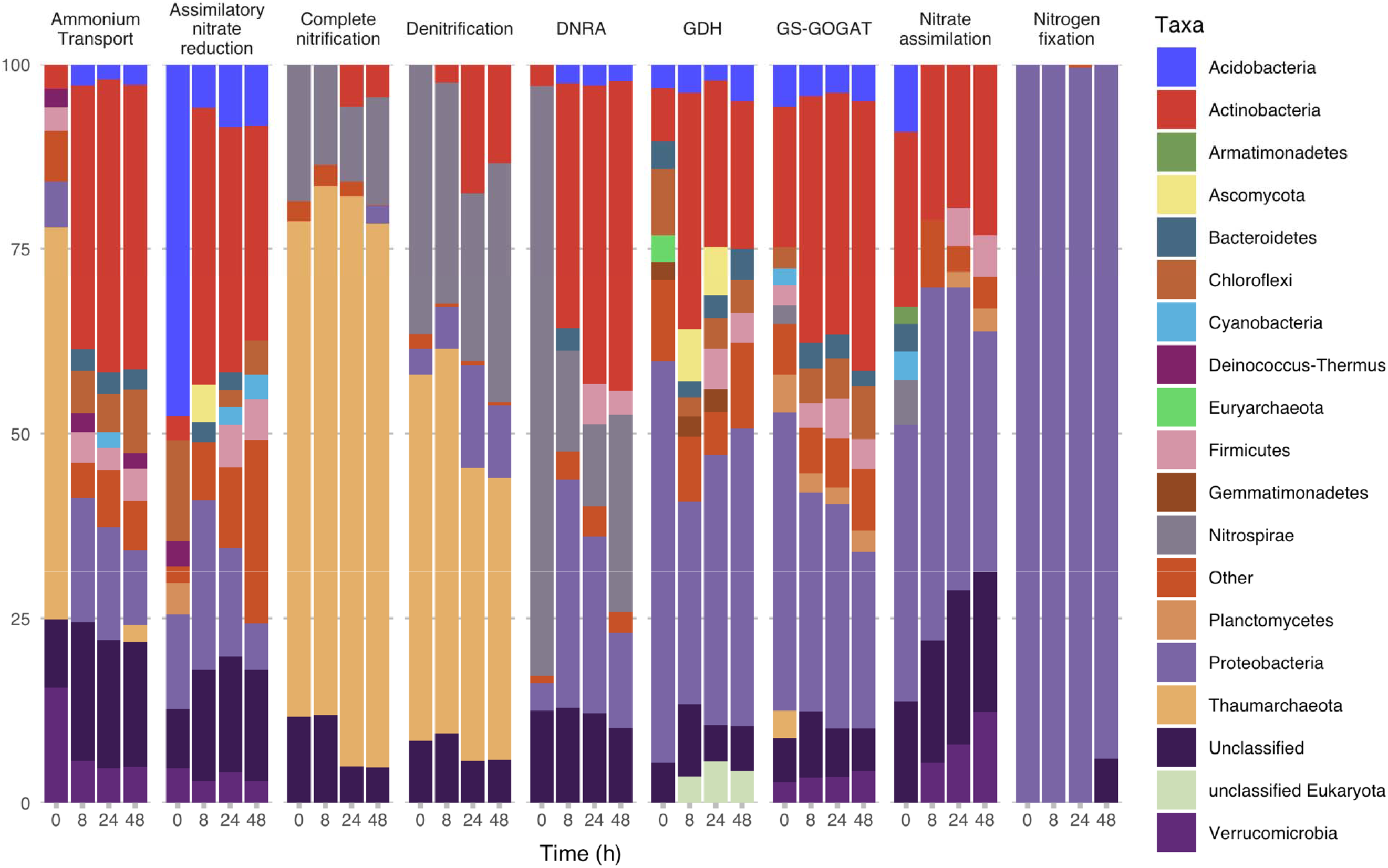
Relative transcript abundance of major taxa for reactions and pathways of N-cycling at 0, 8, 24, and 48 hours after glucose amendments.

### Regulation of N Cycling Genes

Generally, transcripts of genes associated with regulation of N metabolism increased after glucose addition (Fig. S3; Fig. 5). The abundance of ATase and UTase (*glnD* and *glnE)*, used for post-modification of glutamine synthetase (GS) and regulatory PII proteins respectively, initially increased at t_8_ (2.18 ± 0.41 LFC and 4.31 ± 0.36 LFC; FDR < 0.01; Fig. S3; Fig. 5). UTase (*glnD*) but not ATase (glnE), continued to be significantly upregulated at t_24_ (3.79 ± 0.36 LFC) and t_48_ (2.75 ± 0.36 LFC; Fig. S3). Similar upregulation was noted for PII proteins GlnB (*glnB*; LFC > 2.9; FDR < 0.01; Fig. S3) and GlnK (*glnK*; LFC > 3.9; FDR < 0.01; Fig. S3), and the NtrC family genes *glnL* (FDR < 0.01) and *glnG* (FDR < 0.01 at t_8_ and t_24_; Fig. S3). No significant changes in transcript abundances were found for the transcriptional regulators *nac and lrp,* while *crp* and *rpoN* were slightly downregulated (LFC < −1) at t_8_ and t_24_ (FDR < 0.01; Fig. S3; Fig. 5).

**Figure 5.**
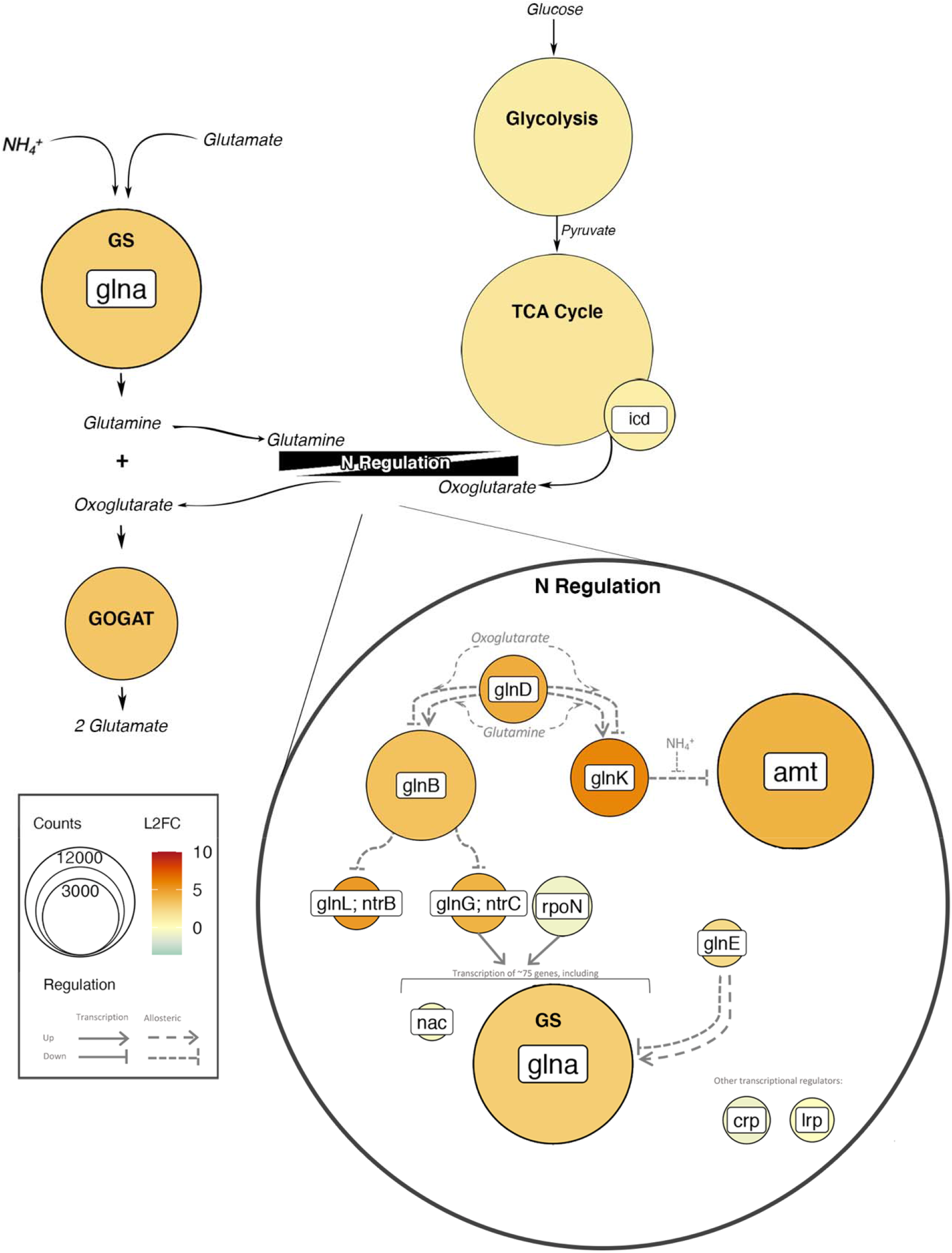
Abundance and log_2_-fold change of transcripts 8 h after glucose addition of C and N metabolism including glycolysis, the TCA cycle, N regulatory network, and GS-GOGAT. Color represents log_2_-fold change of transcript abundances relative to t_0_, and size indicates number of transcripts. Thin black arrows indicate reactants or products of pathways and grey arrows represent regulatory controls. Gene names are presented in white boxes (ex. *glnA*), whereas pathway or enzyme names are presented in bold (ex. GS or Glycolysis).

### C Metabolism

The LFC and total number of normalized transcripts for processes involved with glucose breakdown (KEGG modules M00001, M00003, M00004, M0008, and M00009). increased from t_0_ to t_8_ and t_24_ (Fig. S4; Table S4; Tukey’s HSD p < 0.05). Significant changes in transcript abundance after glucose amendment were found for the Entner-Doudoroff pathway and TCA cycle, including the enzyme isocitrate dehydrogenase (*icd*) which produces oxoglutarate, a metabolite which directly connects C and N metabolism (Fig. 5; Fig. S4B, Table S4).

## DISCUSSION

Over a period of 48-hours after glucose addition we observed a substantial decrease in K_2_SO_4_ extractable organic C, an increase in CO_2_ production rate, and an increase in the abundance of transcripts for genes associated with glucose breakdown. These changes coincided with a decrease in inorganic N and an increase in the transcript abundance of genes involved with inorganic N uptake, assimilation, and N metabolism regulation. These results demonstrate that soil microbial communities respond to labile C not only by upregulating genes associated with C metabolism, but also by rapidly increasing the transcription of genes responsible for N acquisition. Further, we found that genes for several forms of N acquisition (e.g., N fixation, assimilatory nitrate reduction, ammonium transport) were differentially transcribed over the 48 h incubation, indicating that changes in multiple microbially mediated N transformations occur within this small temporal window.

### Inorganic N Uptake and Assimilation

The GS-GOGAT pathway appeared to be the predominant pathway through which ammonium was assimilated into biomass. The other main avenue of ammonium assimilation into biomass, the enzyme GDH, did not show a similar increase in transcript abundance and the abundance of GDH transcripts was substantially smaller than that of GS-GOGAT. This suggests that GS-GOGAT may be the dominant pathway for assimilation of inorganic N in soil microbial communities responding to labile C inputs. This finding is consistent with the notion that GDH is most active when NH_4_^+^ concentration is high and availability of C is low (27). Assays from soil microbial communities have also shown that GS activity increases in response to higher C to N ratios whereas GDH activity decreases (75). Further, we found that regulation of GDH transcription appeared to be gene specific, with transcription for EC 1.4.1.4 increasing early and EC 1.4.1.2 increasing late. These results nicely follow concentrations of NH_4_^+^ as NADPH specific enzymes (EC 1.4.1.4) are generally used for ammonium assimilation (76) whereas NADH specific enzymes (EC 1.4.1.2) are commonly used for breakdown of glutamate to ammonium (77). These findings highlight the potential utility of measuring GDH and GS-GOGAT gene transcription for tracking the C and N balance within microbial communities at a given moment in time, which could be a useful approach when, for example, assessing how specific land use practices influence microbial metabolism and N cycling.

Various mechanisms for transporting inorganic N across the cell membrane were upregulated in response to glucose inputs. Notably, the gene *amtB*, which encodes for the ammonium transporter AmtB, was the second most abundant upregulated gene during the incubation (behind *glnA*). Similarly, we observed an upregulation of genes associated with nitrate and nitrite transport (KEGG module M00615) and assimilatory nitrate reduction, which coincided with a precipitous drop in the concentration of NO_3_^−^. Most genes involved with DNRA were not differentially expressed, indicating that nitrate reduction was primarily occurring under aerobic conditions. A notable exception were the genes *nirB* and *nirD*, which encode for the cytosolic enzyme nitrite reductase NirBD (78), which has been shown to be active in aerobic soils (79, 80) and may function as the nitrite reductase in assimilatory nitrate reduction (81). Although the upregulation of N transport genes in response to glucose is certainly not novel (30), these results are the first demonstration of this response in a soil microbial community metatranscriptome. Further, these responses show the short timeframes (within 8 h) in which soil microbial communities can respond to changes in C and N availability.

The finding that glucose addition strongly upregulated genes encoding for nitrogenase, especially when NH_4_^+^ concentrations were low, is consistent with the idea that nitrogen fixation increases when N concentrations are low (82). N fixation has been shown to be activated by the addition of other limiting nutrients such as carbon or phosphorous (83, 84). We therefore believe that the upregulation of nitrogenase genes is a response to low concentrations of NH_4_^+^ and availability of labile C. The prompt upregulation, and subsequent downregulation, of nitrogenase genes also suggests that some portion of biological nitrogen fixation occurs rapidly in soils, or at the very least that the process is highly sensitive to concentrations of NH_4_^+^.

### Connections Between C and N Metabolism

Interestingly, transcripts associated with NH_4_^+^ and NO_3_^−^ transport maintained their high abundances despite concentrations of NO_3_^−^ stabilizing and concentrations of NH_4_increasing (24–48 h into the incubation). One possible explanation is that the activity of these proteins is dictated through allosteric regulation which is tightly connected to the activity of both C and N metabolism (Fig. 5). For example, the ammonium transporter AmtB is allosterically inhibited by the PII protein GlnK which is indirectly controlled by internal concentrations of glutamine, an intermediate of N uptake through GS-GOGAT (Fig. 5), and oxoglutarate, an intermediate of the TCA cycle (Fig. 5; 29, 66). In this way, internal concentrations of metabolites from both C and N metabolism may dictate N uptake.

The transcription of N regulatory genes reflects the importance of intermediate metabolites in regulation. We found that abundance of transcripts for transcriptional regulators (such as *nac, lrp,* and *crp*) and σ^54^ were either not affected or slightly reduced (Fig. 5). In contrast, transcripts for genes in the phosphorylation cascade, which links C and N metabolism through intermediate metabolites, were more abundant after the addition of glucose (Fig. 5). The upregulation of the two component regulatory NtrB (*glnL*, *ntrB*) and NtrC (*glnG*, *ntrC*) within this cascade is especially noteworthy, as this system regulates ~75 genes associated with N acquisition, including glutamine synthetase (Fig. 5) (86).

Since the activity of this regulatory network is tightly controlled by internal concentrations of metabolites (30), it is not possible to determine the activity of many of these proteins through the metatranscriptome alone. However, it is noteworthy that almost all of the genes within this regulatory network were upregulated, even if the encoded protein potentially inhibited N transport or assimilation (e.g. GlnK; Fig. 5). This broad upregulation of genes in the phosphorylation cascade may be beneficial during C uptake, as it allows the concentration of nutrients and metabolites to control N uptake, thereby ensuring N uptake matches the supply of C (25, 32).

### Nitrification and Denitrification

Most genes associated with nitrification and denitrification were significantly downregulated. Since nearly all nitrifiers in this soil were autotrophic archaea (55), this finding is consistent with the premise that addition of glucose reduces rates of autotrophic nitrification by reducing the amount of available ammonium (37). It is not especially surprising that we did not find an upregulation of denitrification genes, as denitrification is most prevalent in anoxic systems with high availabilities of nitrate.

### Genetic Potential Versus Transcription

Notably, although we did observe a slight shift in the functional composition of our metagenomes, these changes did not track those found in the metatranscriptomes in either magnitude or direction. Changes contributing the most to dissimilarity tended to be slight shifts in highly abundant genes, such as *rpoB*, *rpoC*, and *lacI*. We found interesting differences in the abundance of spore forming proteins as nutrient availability declined, however since many of these proteins were uncommon and in low abundance, the chance of obtaining a false positive is much greater and we are therefore cautious to draw any conclusions based on these data alone. Changes in gene abundance for most N cycling genes were absent. These results suggest that understanding the response of soil microbial communities to short-term changes in the environment necessitates looking beyond the metagenome, as consequential microbial responses occur through changes in gene-expression. This is in line with other studies where the composition of transcripts shifts over hours or days (12, 87), whereas shifts in metagenomic community composition have been shown to occur after weeks or months (72).

Our work represents a preliminary look into the short-term transcriptional response of microbial communities in response to a change in C availability, however there are a number of considerations moving forward. More work needs to be done focusing on this response in a variety soils, as nutrient availability and other soil properties will undoubtably influence this process. For example, soils high in C and low in N would likely not demonstrate a similar response as observed for this agricultural soil. Understanding how ecosystem properties influence the dynamics of transcriptional profiles is therefore necessary in determining short-term microbial contributions to biogeochemical cycling. Further, this work focused on a relatively short timeframe, however whether this increase in transcription persists or influences nutrient cycling on the scale of weeks to months remains to be seen. Finally, future efforts should be made to observe these short-term effects *in situ*. Laboratory incubations are extremely useful for controlling environmental variables and isolating a particular response. However, it is likely that under field conditions, and in the presence of plant roots, factors other than C availability will affect the gene-expression at the same time and to different degrees, potential masking the response observed in this short-term laboratory experiment.

## CONCLUSIONS

Our results indicate strong and rapid upregulation of genes associated with uptake of inorganic N, assimilatory nitrate and nitrite reduction, GS-GOGAT pathway, and the regulatory network underlying N cycling. Further, the majority of upregulation occurred in pathways which are largely aerobic and heterotrophic, suggesting that these processes dominate the short-term response to labile C in these soils. Perhaps most importantly, this work highlights the importance of microbial gene transcription in controlling short-term biogeochemical cycling in soils. Within the 48 h incubation we found that microbially mediated transformations of N were well reflected in the metatranscriptome but not in the metagenome or in microbial biomass. The short-term transcriptional responses of soil microbes may therefore serve an important role in determining how biogeochemical fluxes respond to immediate changes in the environment.

## Supporting information

Supplemental figures and tanles

## ACKNOWLEDGEMENTS

This work was supported by funding from the USDA National Institute of Food and Agriculture Foundational Program (award #2017-67019-26396) and additional support for PD was provided by the U.S. Department of Energy, Office of Biological and Environmental Research, Genomic Science Program LLNL ‘Microbes Persist’ Soil Microbiome Scientific Focus Area (award #SCW1632). The work conducted by the U.S. Department of Energy Joint Genome Institute, a DOE Office of Science User Facility, is supported under Contract No. DE-AC02-05CH11231.

We would like to thank Rebecca Mau, Michaela Hayer, Alicia Purcell, and Ayla Martinez for their assistance with laboratory analyses; Sam Bunkers, Kieston Guidry, and Kiara Nelson for their help downloading and cleaning the data; and Isaac Shaffer for his assistance with the analysis. We would also like to thank the Joint Genome Institute for their work in sequencing and assembly, specifically: Marcel Huntemann, Alicia Clum, Brian Foster, Bryce Foster, Simon Roux, Krishnaveni Palaniappan, Neha Varghese, Supratim Mukherjee, T.B.K. Reddy, Chris Daum, Alex Copeland, Natalia N. Ivanova, Nikos C. Kyrpides, Tijana Glavina del Rio, and Emiley A. Eloe-Fadrosh.

## REFERENCES

1. LeBauer DS, Treseder KK. 2008. Nitrogen limitation of net primary productivity in terrestrial ecosystems is globally distributed. Ecology 89:371–379.

2. Skiba U, Smith KA. 2000. The control of nitrous oxide emissions from agricultural and natural soils. Chemosph - Glob Chang Sci 2:379–386.

3. Camargo JA, Alonso Á. 2006. Ecological and toxicological effects of inorganic nitrogen pollution in aquatic ecosystems: A global assessment. Environ Int 32:831–849.

4. Mooshammer M, Wanek W, Hämmerle I, Fuchslueger L, Hofhansl F, Knoltsch A, Schnecker J, Takriti M, Watzka M, Wild B, Keiblinger KM, Zechmeister-Boltenstern S, Richter A. 2014. Adjustment of microbial nitrogen use efficiency to carbon:Nitrogen imbalances regulates soil nitrogen cycling. Nat Commun 5:1–7.

5. Hallin S, Jones CM, Schloter M, Philippot L. 2009. Relationship between n-cycling communities and ecosystem functioning in a 50-year-old fertilization experiment. ISME J 3:597–605.

6. Batista MB, Dixon R. 2019. Manipulating nitrogen regulation in diazotrophic bacteria for agronomic benefit. Biochem Soc Trans 47:603–614.

7. Marzluf GA. 1997. Genetic regulation of nitrogen metabolism in the fungi. Microbiol Mol Biol Rev 61:17–32.

8. Reitzer L. 2003. Nitrogen assimilation and global regulation in Escherichia coli. Annu Rev Microbiol 57:155–176.

9. Cebolla A, Palomares AJ. 1994. Genetic regulation of nitrogen fixation in Rhizobium meliloti. Microbiologia 10:371–384.

10. Kuzyakov Y, Blagodatskaya E. 2015. Microbial hotspots and hot moments in soil: Concept & review. Soil Biol Biochem 83:184–199.

11. Albright MBN, Johansen R, Lopez D, Gallegos-Graves LV, Steven B, Kuske CR, Dunbar J. 2018. Short-term transcriptional response of microbial communities to nitrogen fertilization in a pine forest soil. Appl Environ Microbiol 84:e00598–18.

12. León-Sobrino C, Ramond J-B, Maggs-Kölling G, Cowan DA. 2019. Nutrient acquisition, rather than stress response over diel cycles, drives microbial transcription in a hyper-arid Namib Desert soil. Front Microbiol 10:1054.

13. Coskun D, Britto DT, Shi W, Kronzucker HJ. 2017. How plant root exudates shape the nitrogen cycle. Trends Plant Sci 22:661–673.

14. Trap J, Bonkowski M, Plassard C, Villenave C, Blanchart E. 2016. Ecological importance of soil bacterivores for ecosystem functions. Plant Soil 398:1–24.

15. Trubl G, Jang H Bin, Roux S, Emerson JB, Solonenko N, Vik DR, Solden L, Ellenbogen J, Runyon AT, Bolduc B, Woodcroft BJ, Saleska SR, Tyson GW, Wrighton KC, Sullivan MB, Rich VI. 2018. Soil Viruses Are Underexplored Players in Ecosystem Carbon Processing. mSystems 3:1–21.

16. Kuzyakov Y. 2010. Priming effects: Interactions between living and dead organic matter. Soil Biol Biochem 42:1363–1371.

17. Demoling F, Figueroa D, Bååth E. 2007. Comparison of factors limiting bacterial growth in different soils. Soil Biol Biochem 39:2485–2495.

18. Hobbie JE, Hobbie EA. 2013. Microbes in nature are limited by carbon and energy: the starving-survival lifestyle in soil and consequences for estimating microbial rates. Front Microbiol 4:324.

19. Schimel JP, Weintraub MN. 2003. The implications of exoenzyme activity on microbial carbon and nitrogen limitation in soil: a theoretical model. Soil Biol Biochem 35:549–563.

20. Papp K, Hungate BA, Schwartz E. 2019. Glucose triggers strong taxonLJspecific responses in microbial growth and activity: insights from DNA and RNA qSIP. Ecology ecy.2887.

21. Kamble PN, Bååth E. 2014. Induced N-limitation of bacterial growth in soil: Effect of carbon loading and N status in soil. Soil Biol Biochem 74:11–20.

22. Geisseler D, Horwath WR, Joergensen RG, Ludwig B. 2010. Pathways of nitrogen utilization by soil microorganisms - A review. Soil Biol Biochem 42:2058–2067.

23. Yang L, Zhang L, Geisseler D, Wu Z, Gong P, Xue Y, Yu C, Juan Y, Horwath WR. 2016. Available C and N affect the utilization of glycine by soil microorganisms. Geoderma 283:32–38.

24. Geisseler D, Horwath WR. 2008. Regulation of extracellular protease activity in soil in response to different sources and concentrations of nitrogen and carbon. Soil Biol Biochem 40:3040–3048.

25. Chubukov V, Gerosa L, Kochanowski K, Sauer U. 2014. Coordination of microbial metabolism. Nat Rev Microbiol 12:327–340.

26. Yuan J, Doucette CD, Fowler WU, Feng X, Piazza M, Rabitz HA, Wingreen NS, Rabinowitz JD. 2009. Metabolomics◻driven quantitative analysis of ammonia assimilation in *E. coli*. Mol Syst Biol 5:302.

27. Sharkey MA, Engel PC. 2008. Apparent negative co-operativity and substrate inhibition in overexpressed glutamate dehydrogenase from *Escherichia coli*. FEMS Microbiol Lett 281:132–139.

28. Lin JT, Stewart V. 1997. Nitrate assimilation by bacteria. Adv Microb Physiol 39:1–30.

29. Zehr JP, Turner PJ. 2001. Nitrogen fixation: Nitrogenase genes and gene expression. Methods Microbiol 30:271–286.

30. van Heeswijk WC, Westerhoff H V., Boogerd FC. 2013. Nitrogen assimilation in *Escherichia coli*: Putting molecular data into a systems perspective. Microbiol Mol Biol Rev 77:628–695.

31. Merrick MJ. 1993. In a class of its own — the RNA polymerase sigma factor σ;54 (σN). Mol Microbiol 10:903–909.

32. Huergo LF, Dixon R. 2015. The emergence of 2-oxoglutarate as a master regulator metabolite. Microbiol Mol Biol Rev 79:419–35.

33. Stein LY, Klotz MG. 2016. The nitrogen cycle. Curr Biol 26:R94–R98.

34. Daims H, Lebedeva E V., Pjevac P, Han P, Herbold C, Albertsen M, Jehmlich N, Palatinszky M, Vierheilig J, Bulaev A, Kirkegaard RH, Von Bergen M, Rattei T, Bendinger B, Nielsen PH, Wagner M. 2015. Complete nitrification by Nitrospira bacteria. Nature 528:504–509.

35. Van Kessel MAHJ, Speth DR, Albertsen M, Nielsen PH, Op Den Camp HJM, Kartal B, Jetten MSM, Lücker S. 2015. Complete nitrification by a single microorganism. Nature 528:555–559.

36. Hu H-W, Chen D, He J-Z. 2015. Microbial regulation of terrestrial nitrous oxide formation: understanding the biological pathways for prediction of emission rates. FEMS Microbiol Rev 021:729–749.

37. Verhagen FJM, Duyts H, Laanbroek HJ. 1992. Competition for ammonium between nitrifying and heterotrophic bacteria in continuously percolated soil columns. Appl Environ Microbiol 58:3303–3311.

38. Lan T, Liu R, Suter H, Deng O, Gao X, Luo L, Yuan S, Wang C, Chen D. 2020. Stimulation of heterotrophic nitrification and N2O production, inhibition of autotrophic nitrification in soil by adding readily degradable carbon. J Soils Sediments 20:81–90.

39. Tiedje JM, Sexstone AJ, Myrold DD, Robinson JA. 1983. Denitrification: ecological niches, competition and survival. Antonie Van Leeuwenhoek 48:569–583.

40. Henderson SL, Dandie CE, Patten CL, Zebarth BJ, Burton DL, Trevors JT, Goyer C. 2010. Changes in denitrifier abundance, denitrification gene mRNA levels, nitrous oxide emissions, and denitrification in anoxic soil microcosms amended with glucose and plant residues. Appl Environ Microbiol 76:2155–2164.

41. Carvalhais LC, Dennis PG, Tyson GW, Schenk PM. 2012. Application of metatranscriptomics to soil environments. J Microbiol Methods 91:246–251.

42. Moran MA. 2009. Metatranscriptomics: Eavesdropping on complex microbial communities. Microbe 4:329.

43. Helbling DE, Ackermann M, Fenner K, Kohler HPE, Johnson DR. 2012. The activity level of a microbial community function can be predicted from its metatranscriptome. ISME J 6:902–904.

44. Nacke H, Fischer C, Thürmer A, Meinicke P, Daniel R. 2014. Land use type significantly affects microbial gene transcription in soil. Microb Ecol 67:919–930.

45. Damon C, Lehembre F, Oger-Desfeux C, Luis P, Ranger J, Fraissinet-Tachet L, Marmeisse R. 2012. Metatranscriptomics reveals the diversity of genes expressed by eukaryotes in forest soils. PLoS One 7:e28967.

46. Žifčáková L, Větrovský T, Howe A, Baldrian P. 2016. Microbial activity in forest soil reflects the changes in ecosystem properties between summer and winter. Environ Microbiol 18:288–301.

47. Kim Y, Liesack W. 2015. Differential assemblage of functional units in paddy soil microbiomes. PLoS One 10:e0122221.

48. Bei Q, Moser G, Wu X, Müller C, Liesack W. 2019. Metatranscriptomics reveals climate change effects on the rhizosphere microbiomes in European grassland. Soil Biol Biochem 138:107604.

49. Baldrian P, Kolařík M, Štursová M, Kopecký J, Valášková V, Větrovský T, Žifčáková L, Šnajdr J, Rídl J, Vlček Č, Voříšková J. 2012. Active and total microbial communities in forest soil are largely different and highly stratified during decomposition. ISME J 6:248–258.

50. Anderson TH, Domsch KH. 1985. Maintenance carbon requirements of actively-metabolizing microbial populations under in situ conditions. Soil Biol Biochem 17:197–203.

51. Van Hees PAW, Jones DL, Finlay R, Godbold DL, Lundström US. 2005. The carbon we do not see - The impact of low molecular weight compounds on carbon dynamics and respiration in forest soils: A review. Soil Biol Biochem 37:1–13.

52. Reischke S, Rousk J, Bååth E. 2014. The effects of glucose loading rates on bacterial and fungal growth insoil. Soil Biol Biochem 70:88–95.

53. Reischke S, Kumar MGK, Bååth E. 2015. Threshold concentration of glucose for bacterial growth in soil. Soil Biol Biochem 80:218–223.

54. Pena-Yewtukhiw EM, Romano EL, Waterland NL, Grove JH. 2017. Soil health indicators during transition from row crops to grass–legume sod. Soil Sci Soc Am J 0:0.

55. Walkup J, Freedman Z, Kotcon J, Morrissey EM. 2020. Pasture in crop rotations influences microbial biodiversity and function reducing the potential for nitrogen loss from compost. Agric Ecosyst Environ 304.

56. Birch HF. 1958. The effect of soil drying on humus decomposition and nitrogen availability. Plant Soil 10:9–31.

57. Barnard RL, Blazewicz SJ, Firestone MK. 2020. Citation Classic Rewetting of soil: Revisiting the origin of soil CO 2 emissions. Soil Biol Biochem 147:107819.

58. Brookes PC, Landman A, Pruden G, Jenkinson DS. 1985. Chloroform fumigation and the release of soil nitrogen: A rapid direct extraction method to measure microbial biomass nitrogen in soil. Soil Biol Biochem 17:837–842.

59. Dijkstra P, Dalder JJ, Selmants PC, Hart SC, Koch GW, Schwartz E, Hungate BA. 2011. Modeling soil metabolic processes using isotopologue pairs of position-specific 13C-labeled glucose and pyruvate. Soil Biol Biochem 43:1848–1857.

60. Nordberg H, Cantor M, Dusheyko S, Hua S, Poliakov A, Shabalov I, Smirnova T, Grigoriev I V., Dubchak I. 2014. The genome portal of the Department of Energy Joint Genome Institute: 2014 updates. Nucleic Acids Res 42.

61. Chuckran PF, Huntemann M, Clum A, Foster B, Foster B, Roux S, Palaniappan K, Varghese N, Mukherjee S, Reddy TBK, Daum C, Copeland A, Ivanova NN, Kyrpides NC, del Rio TG, Eloe-Fadrosh EA, Morrissey EM, Schwartz E, Fofanov V, Hungate B, Dijkstra P. 2020. Metagenomes and Metatranscriptomes of a Glucose-Amended Agricultural Soil. Microbiol Resour Announc 9.

62. Li D, Liu CM, Luo R, Sadakane K, Lam TW. 2015. MEGAHIT: An ultra-fast single-node solution for large and complex metagenomics assembly via succinct de Bruijn graph. Bioinformatics 31:1674–1676.

63. Bankevich A, Nurk S, Antipov D, Gurevich AA, Dvorkin M, Kulikov AS, Lesin VM, Nikolenko SI, Pham S, Prjibelski AD, Pyshkin A V., Sirotkin A V., Vyahhi N, Tesler G, Alekseyev MA, Pevzner PA. 2012. SPAdes: A new genome assembly algorithm and its applications to single-cell sequencing. J Comput Biol 19:455–477.

64. Chen I-MA, Chu K, Palaniappan K, Pillay M, Ratner A, Huang J, Huntemann M, Varghese N, White JR, Seshadri R, Smirnova T, Kirton E, Jungbluth SP, Woyke T, Eloe-Fadrosh EA, Ivanova NN, Kyrpides NC. 2019. IMG/M v.5.0: an integrated data management and comparative analysis system for microbial genomes and microbiomes. Nucleic Acids Res 47:D666–D677.

65. Kanehisa M, Goto S. 2000. Yeast Biochemical Pathways. KEGG: Kyoto encyclopedia of genes and genomes. Nucleic Acids Res 28:27–30.

66. Huber W, Carey VJ, Gentleman R, Anders S, Carlson M, Carvalho BS, Bravo HC, Davis S, Gatto L, Girke T, Gottardo R, Hahne F, Hansen KD, Irizarry RA, Lawrence M, Love MI, MaCdonald J, Obenchain V, Oles◻ AK, Pagès H, Reyes A, Shannon P, Smyth GK, Tenenbaum D, Waldron L, Morgan M. 2015. Orchestrating high-throughput genomic analysis with Bioconductor. Nat Methods 12:115–121.

67. Love MI, Huber W, Anders S. 2014. Moderated estimation of fold change and dispersion for RNA-seq data with DESeq2. Genome Biol 15.

68. Conesa A, Madrigal P, Tarazona S, Gomez-Cabrero D, Cervera A, McPherson A, Szcześniak MW, Gaffney DJ, Elo LL, Zhang X, Mortazavi A. 2016. A survey of best practices for RNA-seq data analysis. Genome Biol 17:13.

69. Oksanen AJ, Blanchet FG, Kindt R, Legen- P, Minchin PR, Hara RBO, Simpson GL, Solymos P, Stevens MHH. 2019. vegan: Community Ecology Package.

70. Kanehisa M, Sato Y. 2019. KEGG Mapper for inferring cellular functions from protein sequences. Protein Sci.

71. Wickham H. 2016. ggplot2: Elegant Graphics for Data Analysisle. Springer-Verlag New York.

72. Team RC. 2018. R: A language and environment for statistical computing. R Found Stat Comput Vienna, Austria.

73. Mooshammer M, Wanek W, Zechmeister-Boltenstern S, Richter A. 2014. Stoichiometric imbalances between terrestrial decomposer communities and their resources: Mechanisms and implications of microbial adaptations to their resources. Front Microbiol 5:22.

74. Smith EL, Austen BM, Blumenthal KM, Nyc JF. 1975. Glutamate DehydrogenasesEnzymes 3rd ed. Academic Press.

75. Geisseler D, Doane TA, Horwath WR. 2009. Determining potential glutamine synthetase and glutamate dehydrogenase activity in soil. Soil Biol Biochem 41:1741–1749.

76. Duncan PA, White BA, Mackie RI. 1992. Purification and properties of NADP-dependent glutamate dehydrogenase from Ruminococcus flavefaciens FD-1. Appl Environ Microbiol 58:4032–4037.

77. Miller SM, Magasanik B. 1990. Role of NAD-linked glutamate dehydrogenase in nitrogen metabolism in Saccharomyces cerevisiae. J Bacteriol 172:4927–4935.

78. Cole J. 1996. Nitrate reduction to ammonia by enteric bacteria: redundancy, or a strategy for survival during oxygen starvation? FEMS Microbiol Lett 136:1–11.

79. Ruiz B, Le Scornet A, Sauviac L, Rémy A, Bruand C, Meilhoc E. 2019. The nitrate assimilatory pathway in Sinorhizobium meliloti: Contribution to NO production. Front Microbiol 10:1526.

80. Pathan SI, Větrovský T, Giagnoni L, Datta R, Baldrian P, Nannipieri P, Renella G. 2018. Microbial expression profiles in the rhizosphere of two maize lines differing in N use efficiency. Plant Soil 433:401–413.

81. Stolz JF, Basu P. 2002. Evolution of nitrate reductase: Molecular and structural variations on a common function. ChemBioChem 3:198–206.

82. Dixon R, Kahn D. 2004. Genetic regulation of biological nitrogen fixation. Nat Rev Microbiol 2:621–631.

83. Benner JW, Vitousek PM. 2007. Development of a diverse epiphyte community in response to phosphorus fertilization. Ecol Lett 10:628–636.

84. Vitousek PM, Menge DNL, Reed SC, Cleveland CC. 2013. Biological nitrogen fixation: Rates, patterns and ecological controls in terrestrial ecosystems. Philos Trans R Soc B Biol Sci 368:1–9.

85. Coutts G. 2002. Membrane sequestration of the signal transduction protein GlnK by the ammonium transporter AmtB. EMBO J 21:536–545.

86. Zimmer DP, Soupene E, Lee HL, Wendisch VF, Khodursky AB, Peter BJ, Bender RA, Kustu S. 2000. Nitrogen regulatory protein C-controlled genes of *Escherichia coli*: Scavenging as a defense against nitrogen limitation. Proc Natl Acad Sci U S A 97:14674–14679.

87. Nuccio EE, Starr E, Karaoz U, Brodie EL, Zhou J, Tringe SG, Malmstrom RR, Woyke T, Banfield JF, Firestone MK, Pett-Ridge J. 2020. Niche differentiation is spatially and temporally regulated in the rhizosphere. ISME J 14:999–1014.

88. Mau RL, Liu CM, Aziz M, Schwartz E, Dijkstra P, Marks JC, Price LB, Keim P, Hungate BA. 2015. Linking soil bacterial biodiversity and soil carbon stability. ISME J 9:1477–1480.

