## Supplemental figures and tanles for "Rapid response of nitrogen cycling gene transcription to labile carbon amendments in a soil microbial community"

SUPPLEMENTAL MATERIAL

Supplemental Figure S1:

Total respiration (µmol CO_2_ g^-1^ soil), over the course of a 48 hour incubation with water and water+glucose.

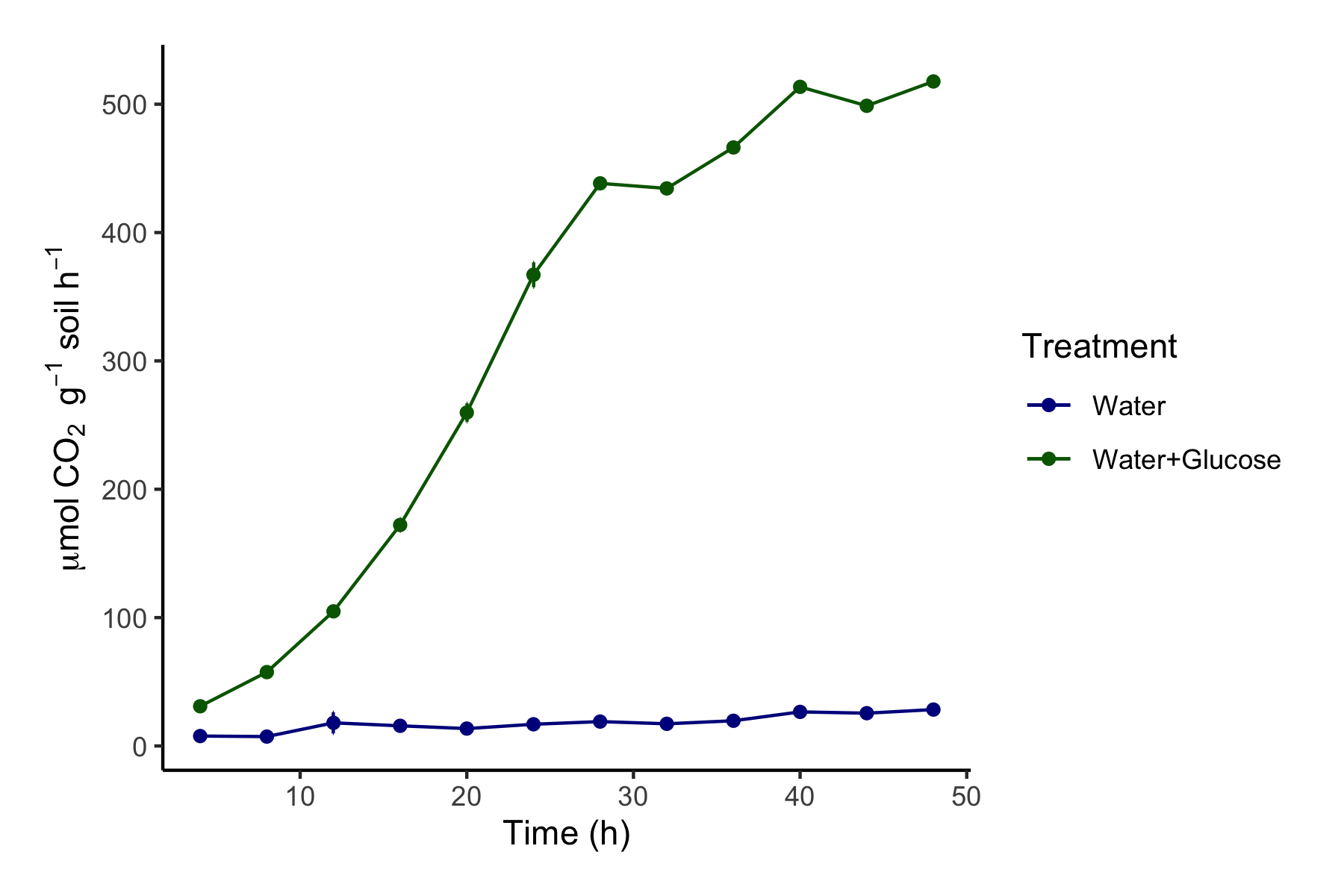

Supplemental Figure S2: Microbial biomass carbon (**A**) and nitrogen (**B**) throughout 48 h incubation after the addition of glucose.

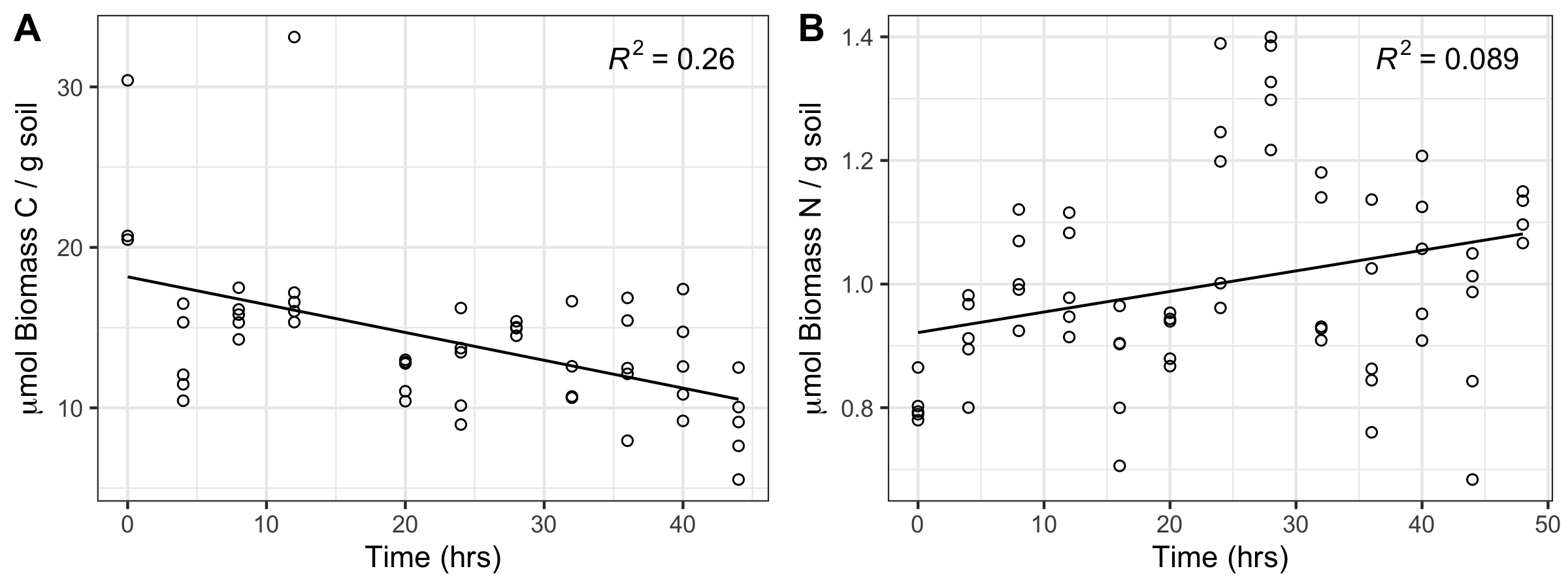

Supplemental Figure S3: Log_2­_-fold change in transcript abundances relative to t_0_ for N regulatory genes for time after glucose amendment (t_8_, t_24,_ and t_48_). Significant changes are indicated with an asterisk.
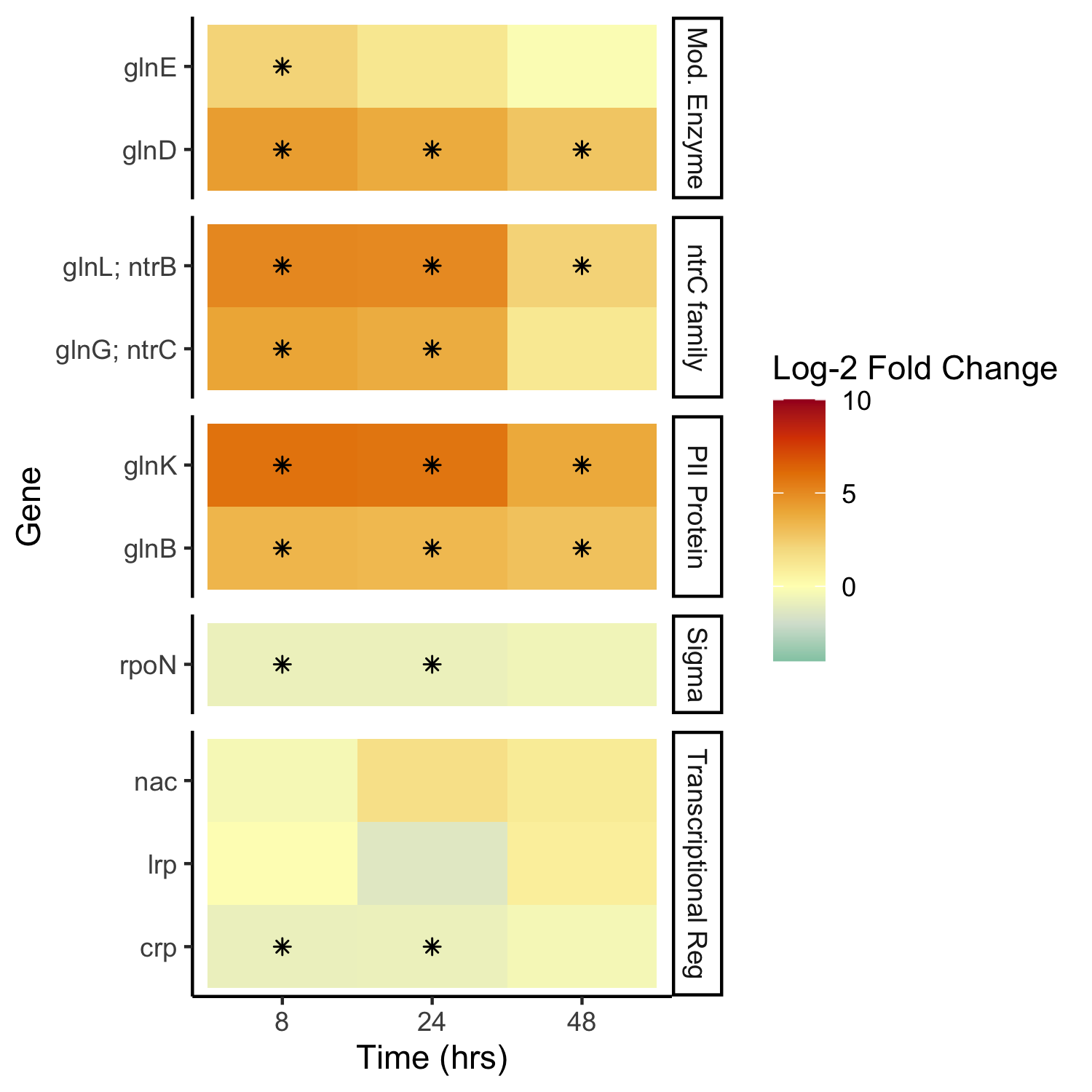

Supplemental Figure S4: The influence of glucose amendments on C metabolism. **(A)** The normalized transcript counts over time for genes associated with glucose catabolism. (**B)** The log_2_-fold change of transcription vs time 0 for each pathway as well as for glycolysis. Significant differences vs time 0 (Tukey’s HSD; P < 0.05) are indicated with an asterisk.

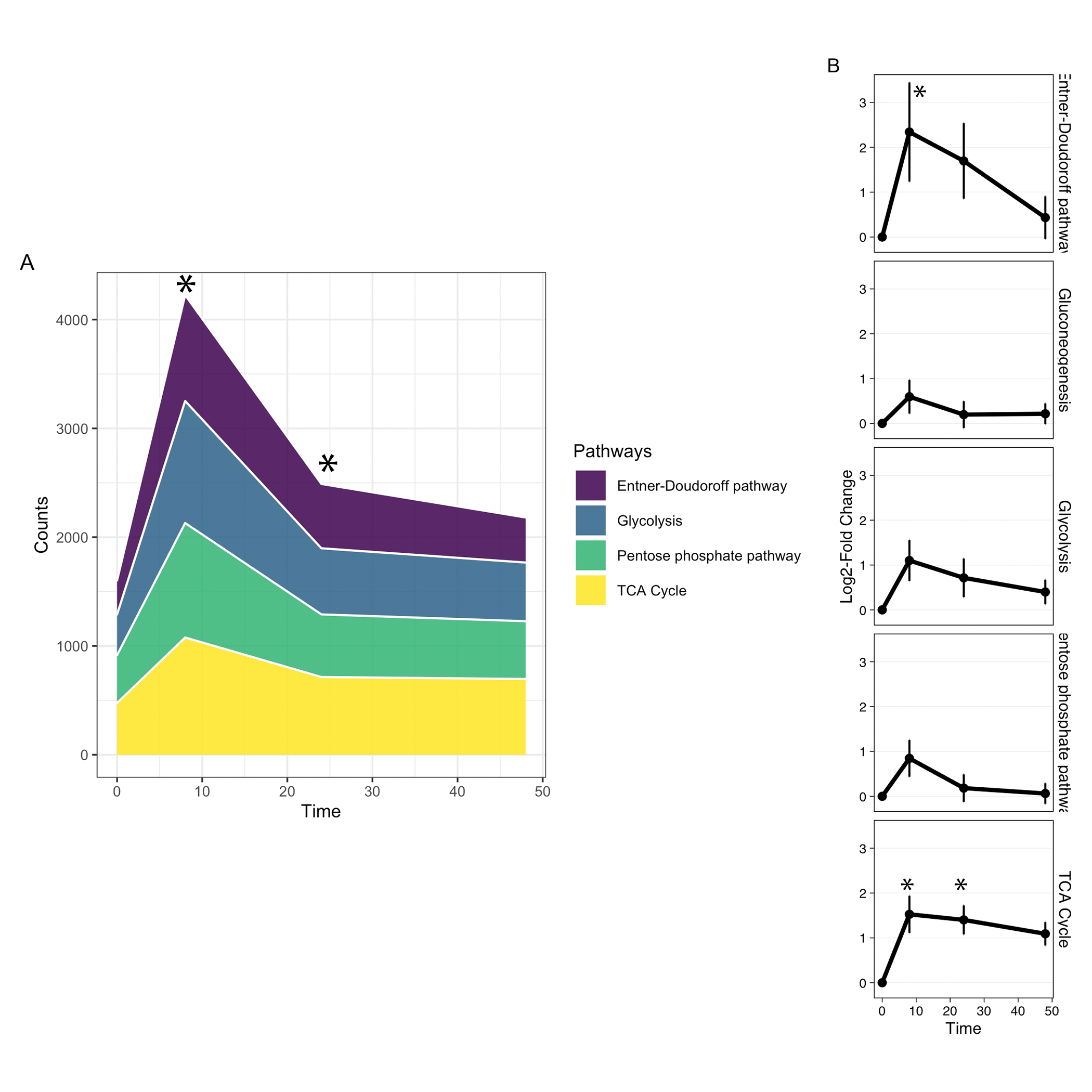

Supplemental Table S1: N cycling genes grouped by reactions and pathways.
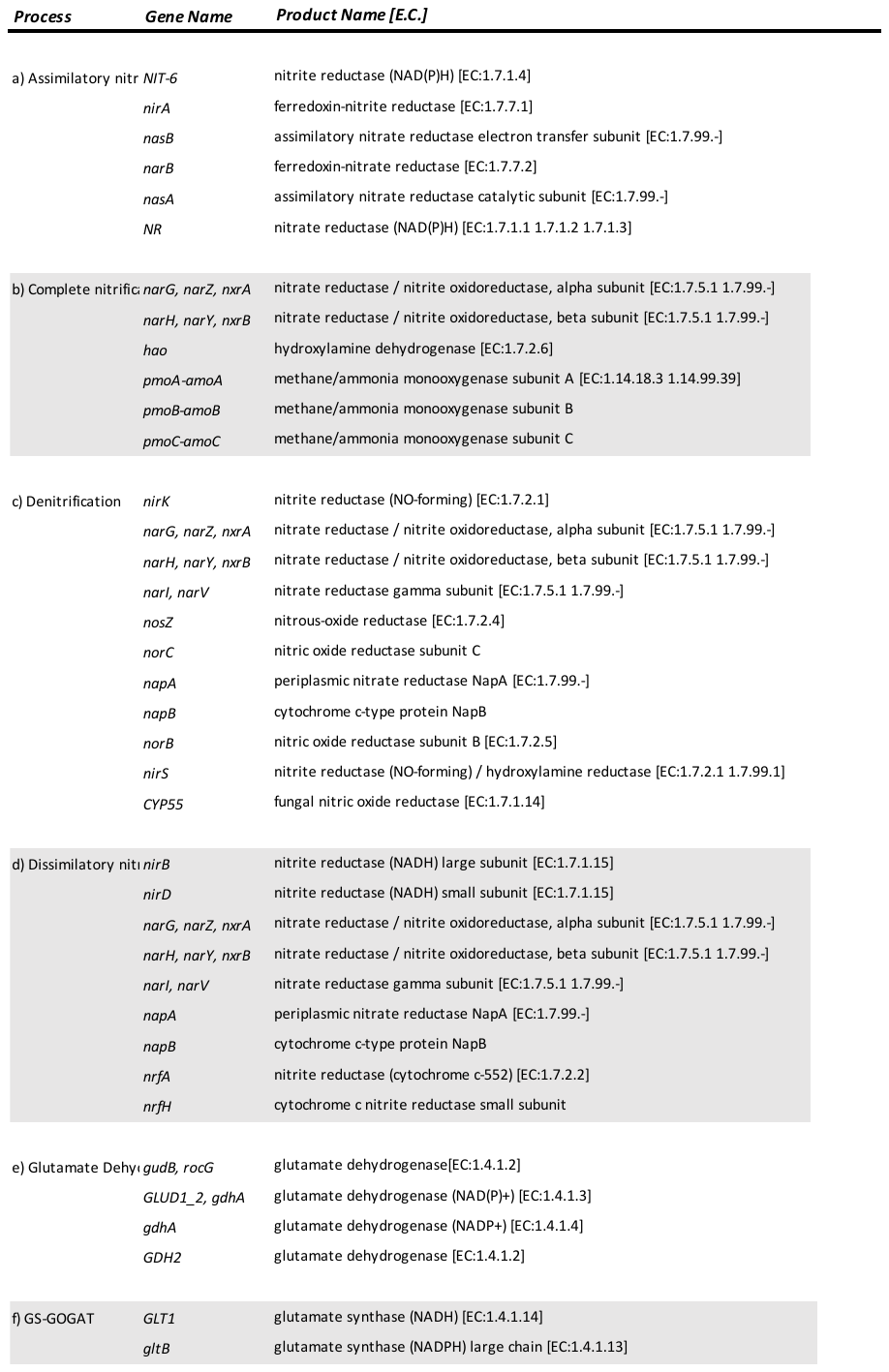

Supplemental Table S2: Differential expression of N cycling genes of soils at 8, 24, and 48 h after glucose amendment vs control. Log_2_-fold change values displayed in bold have a false discovery rate (FDR) of < 0.01, italicized values have a FDR of < 0.1. Base Counts represents the average number of normalized counts across all samples.

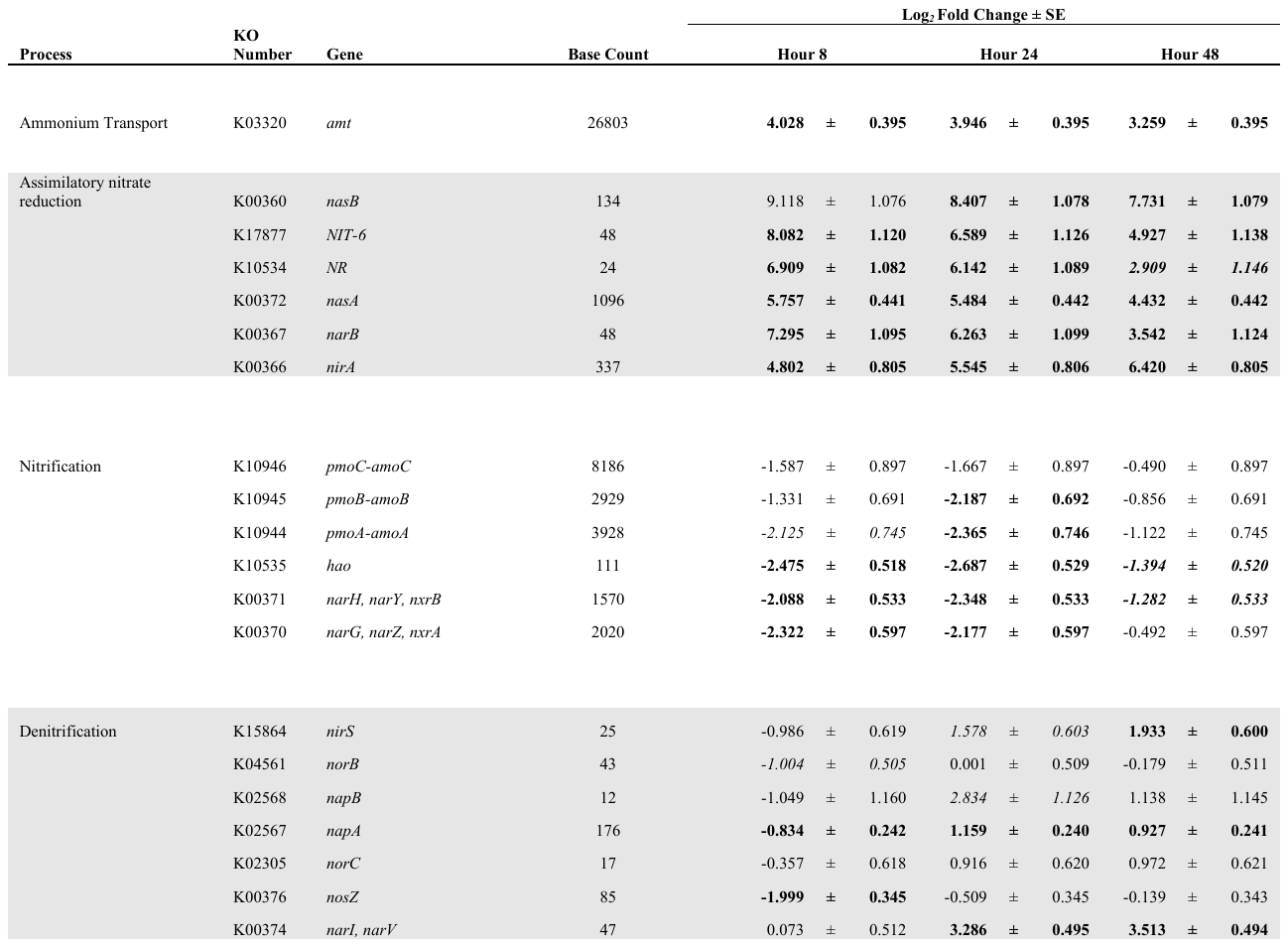

Supplemental Table S3:

Changes in gene abundance in metagenomes. **A)** Results of a SIMPER analysis for the metagenomes 8, 24, and 48 h after addition of glucose amendment, versus the control (t_0_). Shown are the top 3 genes which were most dissimilar in the genetic profile of the treated samples. Values are shown as the percent (%) contribution to dissimilarity of each gene. **B)** Most differentially abundant genes vs t_0_ during the incubation, sorted by LFC.

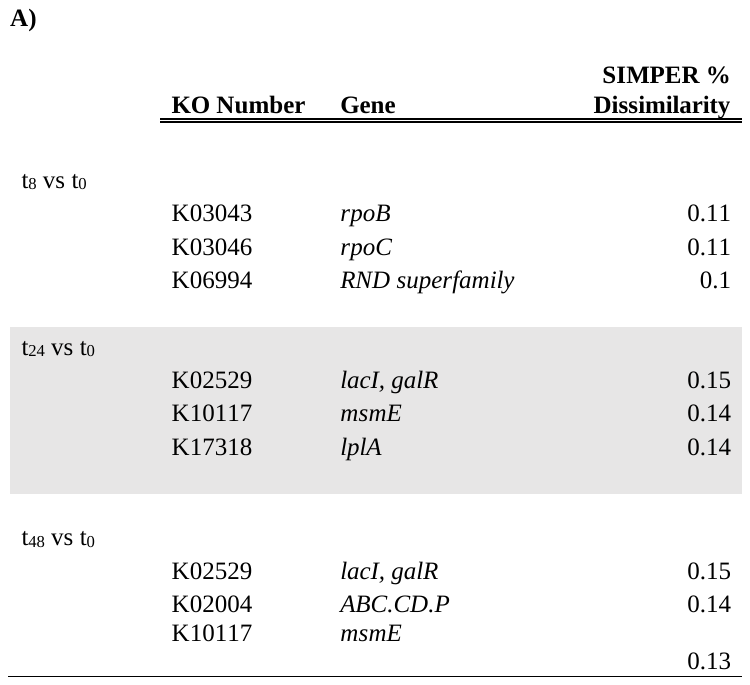

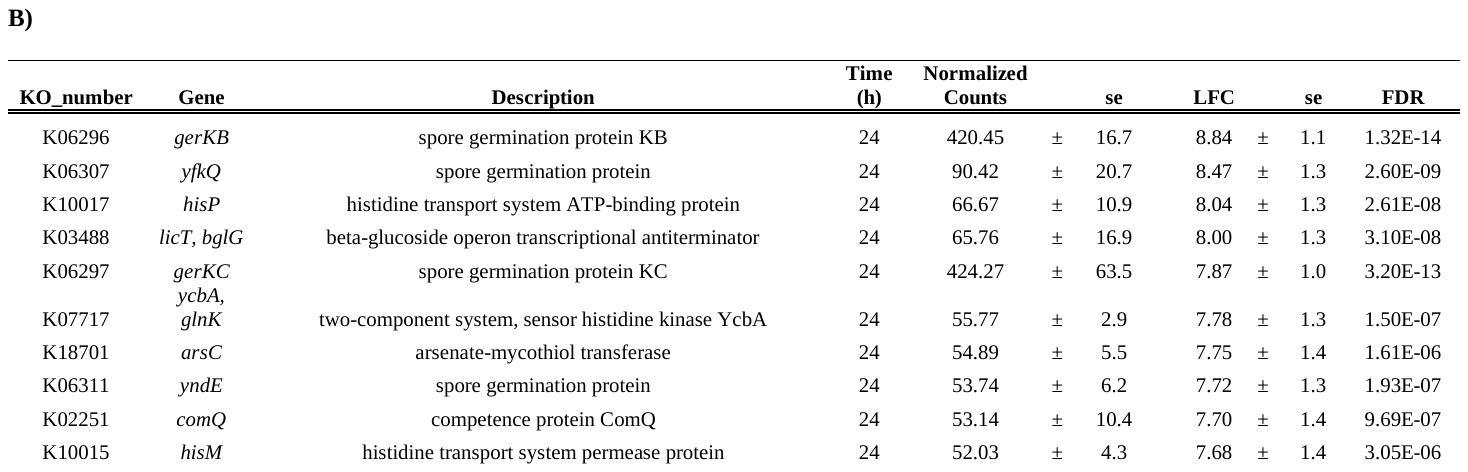
